# Species in big angiosperm genera are priorities for conservation

**DOI:** 10.1101/2025.09.14.674653

**Authors:** Matilda J.M. Brown, Tarciso C.C. Leão, Félix Forest, Eve Lucas, Barnaby E. Walker, Eimear Nic Lughadha

**Affiliations:** Royal Botanic Gardens, Kew, Richmond, Surrey, TW9 3AE, UK

**Author notes:** **Corresponding author:** Matilda Brown, Royal Botanic Gardens, Kew, Richmond, Surrey, TW9 3AE, UK. These two authors contributed equally to this work (Matilda J.M. Brown, Tarciso C.C. Leão).

**Keywords:** Big genera, range size, extinction risk, Red List, angiosperms

## Abstract

Species in big angiosperm genera (≥500 species) are often considered to be less well-known than those in smaller genera. Consequently, species in big genera are likely underrepresented in conservation at all levels, from analyses to actions. However, emerging evidence indicates that species in larger genera tend to have smaller geographic ranges (a strong correlate of increased extinction risk). Here, we test the generality of this pattern by quantifying the relationships between plant taxon size, extinction risk and climatic zone on a global scale. We find that species in larger genera are more likely to have small ranges, less likely to have a global Red List assessment, and, when assessed, are more likely to be threatened. Persistent obstacles to improving conservation documentation of big genera include taxonomic uncertainty, data shortfalls and number of species involved, requiring continued collaboration between taxonomy and conservation to reduce this bias and enable effective conservation of the quarter of plant species that big genera encompass.

**Article Impact Statement:** Species in big angiosperm genera are under-studied, under-assessed, and their extinction risk has been underestimated.

## Introduction

‘Big’ plant genera (≥500 species per Frodin, 2004) contain over a quarter of angiosperm species (Moonlight *et al*., 2024) and include several ecologically, culturally and economically important taxa (e.g. *Euphorbia*, >2,000 spp.; *Solanum*, >1,200 spp., *Eucalyptus*, >700 spp.). Despite the familiarity of many big plant genera, there remains a pervasive knowledge shortfall for species in big genera (Moonlight *et al*., 2024) largely due to the practical challenges associated with working with groups of ≥500 species – through this lens, genus size can be viewed as an attribute that has an effect on species’ individual likelihood of being studied. For example, big genera are less likely to have a global taxonomic monograph and have lower proportions of species with available DNA sequence data (Grace *et al*., 2021). Longstanding latitudinal biases in taxonomic effort (the ‘Latitudinal Taxonomy Gradient’□; Freeman & Pennell, 2021; Guedes *et al*., 2025) mean that this shortfall is particularly severe in the tropics, where the majority of big plant genera occur (Moonlight *et al*., 2024; though this is an effect of overall biodiversity concentration rather than a specific concentration of big genera in the tropics; Baldaszti *et al*., 2025).

We expect that both the practical and taxonomic challenges associated with big plant genera limit our knowledge of extinction risk of species in these taxa; for example, <5% of the >3,000 species of *Astragalus* have global Red List assessments (far fewer than the c. 20% coverage of angiosperms at large; IUCN, 2024), but the effect of genus size on assessment probability has not been quantified. Undertaking a fully-documented species conservation assessment (e.g. a Red List assessment; IUCN, 2012, 2022a) is expensive in terms of time and expertise. Thus, species considered data-poor, which are likely to be assessed as Data Deficient (Brummitt *et al*., 2024) tend to be low priorities for assessment. Similarly, an assessment is specific to a particular species concept, so species that are more likely (or perceived to be more likely) to be re-circumscribed are less likely to be prioritised for assessment. Assessments are typically undertaken as part of projects of clearly defined geographic and/or taxonomic scope, often aiming to assess all species within that scope (e.g. the recent assessment of Dalbergia; Cowell *et al*., 2022; and 332 Mozambican endemics Darbyshire *et al*., 2019). While the global distribution of big genera (Baldaszti *et al*., 2025) means that species in big genera may be proportionally represented in geographically defined projects, they are rarely the taxonomic focus of assessment projects. Comprehensive assessment of species in a big genus requires substantial resources, and usually many collaborators. The successful Global Tree Assessment (>45,000 species; Rivers, 2017) demonstrates the feasibility of large assessment projects, but also the enormity of the challenges involved (Newton *et al*., 2015). We expect that the sheer numbers of species and the (real or perceived) risk of taxonomic uncertainty due to e.g. lack of monographs for big genera mean that a) they are systematically underrepresented on the global Red List and b) if assessed, are more likely to be assessed as Data Deficient.

There is also emerging evidence that we may be underestimating extinction risk in big plant genera. Geographic range size is the strongest correlate of plant extinction risk (Nic Lughadha *et al*., 2019; Walker *et al*., 2022; Bachman *et al*., 2024), and both region-specific and flora-based studies have found that taxa with larger numbers of species have, on average, species with smaller geographic ranges (Schwartz & Simberloff, 2001; Lozano & Schwartz, 2005; Leão *et al*., 2020) and higher proportions of threatened species (Fu *et al*., 2022). Furthermore, a disproportionate number of species described in the last two decades – the majority of which are likely to be threatened (Brown *et al*., 2023a) – are in the very largest genera ( >1,000 species; Moonlight *et al*., 2024). The global generality of the relationships between genus size, range size and extinction risk in flowering plants remain untested, particularly across climatic zones, lifeforms and higher taxonomic ranks, but we expect that an overall genus size - range size relationship, if supported, will also result in higher proportions of threatened species in larger genera.

Here, we test the generality of the negative relationship between genus size and species geographic range size across climate, life form and family, using data from the World Checklist of Vascular Plants (‘WCVP’; Govaerts *et al*., 2021). We then quantify the extent to which species in larger genera are more or less likely to have a Red List assessment, and their probability of, when assessed, being categorised as either Data Deficient or threatened with extinction, using data from the IUCN Red List (hereafter ‘Red List’; IUCN, 2024) and comprehensive species-level extinction risk predictions from Bachman *et al*. (2024). Finally, we identify specific priority big genera for conservation assessment as those for which overall extinction risk on the Red List is most underestimated when compared to predictions.

## Materials and Methods

### Data

We calculated genus size as the number of accepted species in each genus in version 13 of the World Checklist of Vascular Plants (Govaerts *et al*., 2021), accessed via the ‘rWCVP’ and ‘rWCVPdata’ R packages (Brown *et al*., 2023b). Most species lack comprehensive and/or unbiased point data from which to measure range size; instead, as a proxy for range size, we used the number of Level 3 Areas (hereafter ‘botanical countries’, following e.g. Bachman *et al*., 2024; Svahnström *et al*., 2025) of the World Geographic Scheme for Recording Plant Distributions (Brummitt, 2001) accessed via WCVP. Although number of botanical countries is an imperfect proxy for range size (skewed towards single-country endemics, low spatial resolution and unequally-sized botanical countries), it remains the most comprehensive and least biased available data for angiosperms. We acknowledge that these data cannot be said to be a direct measure of range size, but instead represent an intermediate proxy, where species limited to a single botanical country are more likely to have smaller ranges than those occurring in several botanical countries (Svahnström *et al*., 2025). While there are specific scenarios where the relationship between range size and number of botanical countries breaks down (e.g. range-restricted species occurring across country borders, species that are widespread within a large botanical country), we assume that at a global scale, these cases are unlikely to be present with enough frequency to significantly affect our results. Empirical support for this assumption has been reported from a range of studies, where results are consistent across multiple proxies of range size (e.g. extent of occurrence; Bureš *et al*., 2024; Svahnström *et al*., 2025).

We extracted lifeform information from WCVP, then simplified the Raunkiaer lifeforms into annual, herbaceous perennial, tree, other woody perennial, and epiphyte following Humphreys et al. (Humphreys *et al*., 2019) with modifications (see Table S1). For climatic zones, we derived the broad climate types (tropical, subtropical, temperate) from the WCVP ‘climate description’ field or species’ distributions; species spanning multiple climate zones were combined into a ‘multiple climates’ category (Supplementary text; Fig. S1). We focused on the genus-level effects of climate and lifeform, rather than species-level (e.g. species in broadly tropical genera). To assign genus-level climate and lifeform categories, we aggregated species-level climate and lifeform data from WCVP to genus-level using a 75% dominance threshold – e.g., 89.3% of species in *Tillandsia* are epiphytic and 81.4% are tropical, so we treated *Tillandsia* as a tropical, epiphytic genus. Where no single lifeform or climate dominated, genera were assigned to a single ‘multiple climates/lifeforms’ category. This aggregated approach had the additional benefit of allowing us to include in our analyses the c. 78,480 species lacking lifeform information in WCVP version 13. We obtained extinction risk assessments from the IUCN Red List of Threatened Species version 2024-1 (IUCN, 2024), and extinction risk predictions from Bachman *et al*. (2024) using ‘rWCVP’ (Brown *et al*., 2023b) for taxonomic reconciliation, including matching of basionyms, where the taxonomic concept matches that of the accepted species name.

### Statistical analysis

We used logistic regression to model five response variables as a function of genus size: (1) range size, (2) probability of having a global Red List assessment, (3) probability of being categorised as Data Deficient on the Red List, (4) probability of being assessed as threatened on the Red List (i.e. categorised as Vulnerable, Endangered or Critically Endangered), and (5) probability of being either assessed as or predicted to be threatened. Genus size was log_10_-transformed prior to analysis; all effect sizes represent changes on this scale. To investigate the consistency of the genus size-range size relationship across angiosperm diversity, we included three covariates in our analyses: climatic zone, lifeform and family. For each of the five response variables fitted, our overall approach was the same. We first fitted a ‘full model’, which included all covariates and their interaction with genus size (model formula given in Eqn 1; + denotes additive terms; * denotes additive and interaction terms).

***Equation 1***. *Response ∼ GenusSize*Climate + GenusSize*Lifeform + GenusSize*Family*

We then performed model selection using AICc (via the ‘dredge’ function implemented in ‘MuMIn’□; Barton, 2024) to identify the best model/s. To interpret the effect sizes and their associated uncertainty for models with multiple categorical interaction terms, we used estimated marginal means and estimated marginal trends for each value of covariates using the ‘emmeans’ and ‘emtrends’ functions from the ‘emmeans’ R package (Lenth, 2025). To account for imbalances in these variables, we weighted the non-focal covariates by their proportional frequency in the dataset (weights = “cells”). For models that included a family-level interaction term (including those used for model selection), we removed families with fewer than five genera or fewer than 500 species. Although this retained only 25% of angiosperm families, the effect on total species and generic diversity was small; the reduced dataset of 100 families included 89.3% of angiosperm genera and 91.8% of angiosperm species. Removing Aquifoliaceae (which comprises one genus), Balsaminaceae (two genera), Begoniaceae (two genera), Dioscoreaceae (four genera) and Ebenaceae (three genera) resulted in the removal of five big genera from our initial (full) models: *Begonia* (2,092 species), *Dioscorea* (635 species), *Diospyros* (778 species), *Ilex* (564 species) and *Impatiens* (1,120 species). For models fitted to assessed species only, we removed families with fewer than 250 assessments on the Red List (49 families). To check for bias resulting from these reductions, we also fitted models without family effects to the full species dataset.

To account for the skew in range size distribution and lack of fine-scale resolution when using botanical countries, we modelled range size as a function of genus size using a series of logistic regressions. We repeated model-fitting at multiple range-size thresholds using a ‘multiple threshold’ approach to test the robustness of this relationship at different scales. We modelled the probability that species are endemic to a single botanical country, restricted to two or fewer botanical countries, then to three, four and ten or fewer botanical countries.

We used the same ‘multiple threshold’ approach to model extinction risk as for previous studies (Brown *et al*., 2023a; Soto Gomez *et al*., 2024); for the full models including climate, lifeform and family we focused on a single threshold: the probability of being ‘threatened’ per the IUCN (Critically Endangered, Endangered, Vulnerable; IUCN, 2022a), or predicted to be threatened with high confidence (Bachman *et al*., 2024). Species assessed as Extinct (84 species in reduced dataset) or Extinct in the Wild (31 species) could not be modelled separately, so were included under Critically Endangered for all analyses.

To highlight the implications of under-assessment of species in big genera, we directly compared the proportion of threatened species (using the threshold above, noting that this differs from the IUCN calculation in the handling of DD species; IUCN, 2022b) across genus sizes based on assessments only, and assessments plus predictions. From these data, we extracted the 20 big genera with the highest mismatch between their proportion threatened per the Red List, and their estimated proportion threatened from the Bachman et al. (2024) predictions. We present these genera as potential priorities for conservation assessment.

All models were fitted with species as the observation (i.e. data were not aggregated to genus prior to analyses), in R version 4.4.1 (R Core Team, 2024). All data, scripts and outputs required to reproduce our results are available at [link removed for double-blind review].

## Results

Nearly half of all big genera (and species therein) are tropical (Fig 1, Table S2). Of the 82 currently accepted big genera (from version 13 of the WCVP), 40 (49%) are entirely or mostly tropical, containing 40,631 species (48% of species in big genera; 13% of total angiosperm diversity). All big epiphytic genera (10 genera; 11,370 species) are predominantly tropical; similarly, all big tree genera are either tropical (4 genera; 3,211 species) or subtropical (*Eucalyptus*, 713 species). By contrast, most of the 15 temperate big genera are dominated by herbaceous perennials (12 genera; 16,239 species). There are no big genera dominated by annual species.

**Figure 1.**
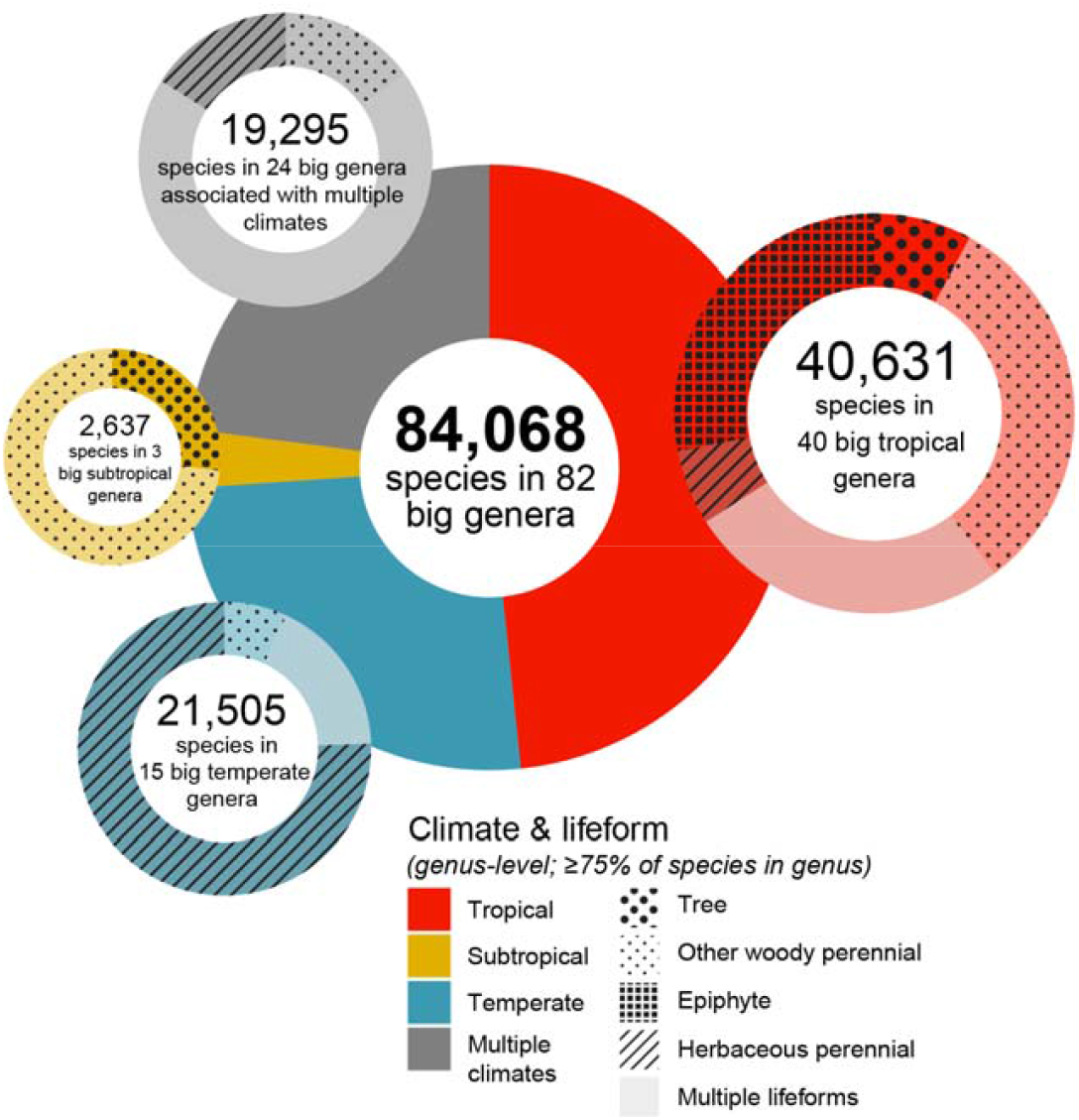
Climate and lifeform distribution in big angiosperm genera. Genera coded by their dominant (≥75% of species) climate/lifeform or as ‘multiple climates/lifeforms’ (see Methods). Colour and pattern show proportion of species across covariate combinations; note that overall circle sizes are not to scale. For full details on numbers of species and genera across covariates, see Table S1.

### Covariates of genus size trends

The best-supported model formula (ranked using AICc; Table S3) across all response variables includes both additive and interaction terms for all covariates (climate, lifeform, family). Family-level interaction terms could not be estimated for families with very few species or genera; the results presented below are thus based on a reduced dataset (91.8% of angiosperm species; see Methods). Models fitted using the full dataset (excluding family effects) are consistent with the findings reported below (Tables S4-6).

### Genus size and species range size

Overall, species in big genera tend to occur in fewer botanical countries so are, on average, more likely to have smaller ranges (Fig 2). The probability of being restricted to a single botanical country is 49.7% (95% CI: 49.2-50.2%) for a species in a monotypic genus, compared to 60.6% (CI: 60.3-60.9%) for species in the ‘smallest’ big genera (i.e. genera of exactly 500 species; from marginal mean estimate; Fig 2a).

**Figure 2.**
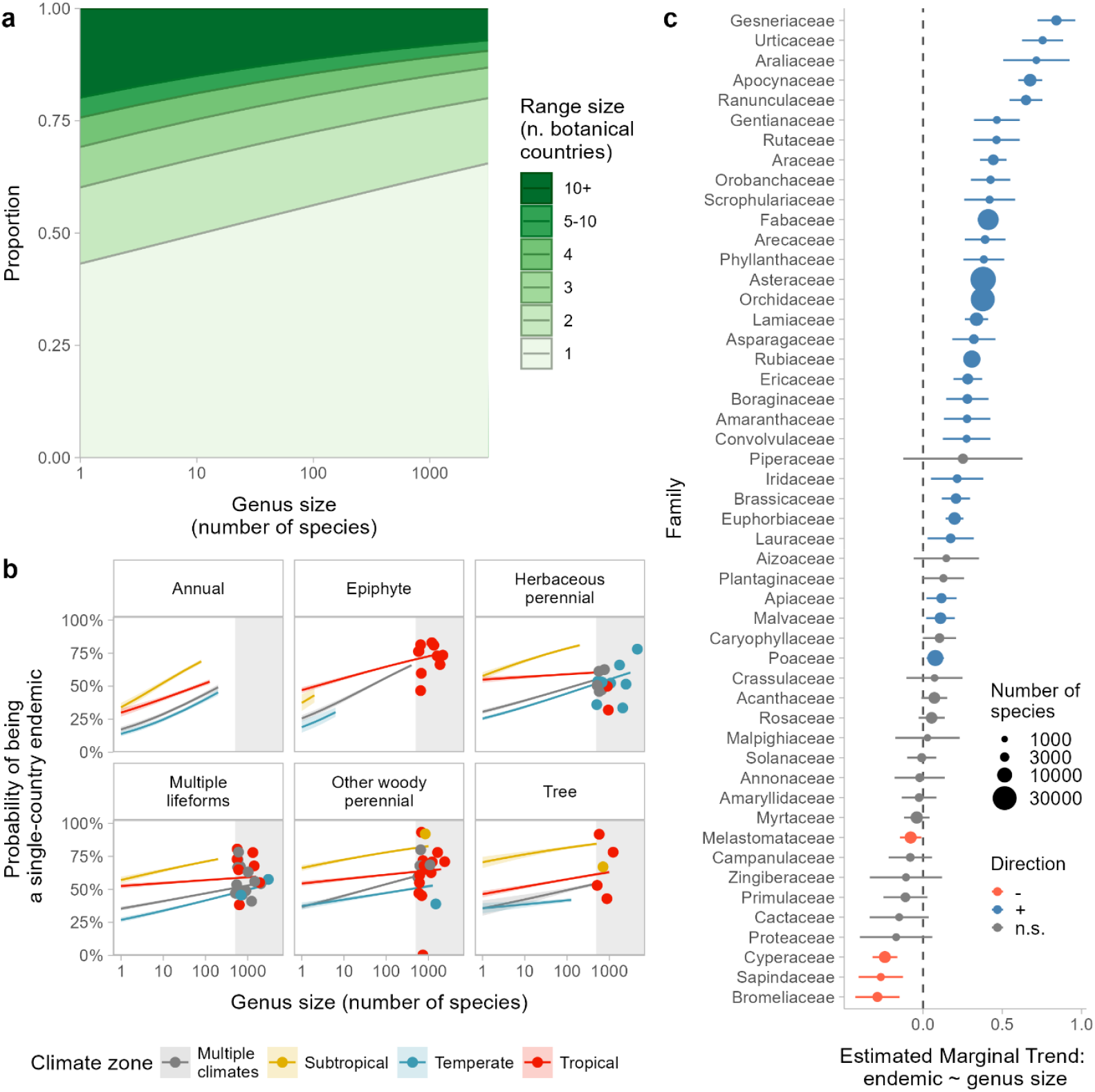
Genus size-range size relationship. Panel a) shows overall estimated marginal mean proportions of species under each range size threshold (as number of botanical countries) across genus size. The median range size is one botanical country for all but the very smallest genera. Panel b) shows the estimated marginal mean probability that species are restricted to a single botanical country across lifeform and climate (note that climate and lifeform aggregated at genus level). Shaded grey area shows the ‘zone of big genera’; points show the observed proportions of single-country endemics in big genera. Panel c) shows the estimated marginal slope coefficients for family (the single-country endemic threshold; only largest 50 families shown); positive coefficients indicate that species in larger genera are more likely to be restricted to a single botanical country. Units of coefficients on the logit scale. Error bars show 95% CIs of coefficient estimates; colour of points denotes significance and direction; size of points denotes number of species per family. Marginal means and trends estimated across cell weights of the non-focal terms in all panels; see Fig S1 for a graphical representation of how these are estimated from the model. For tabulated results with 95% CI values for marginal climate, lifeform, and family trends, see Table S4; for all trends (not averaged), see Table S5.

The relationship between genus size and range size is consistent across scales (i.e. when modelling the proportion of species restricted to one, two, three, four, five and ten botanical countries, Fig 2a), climatic zones and lifeform (Fig 2b). It is strongest for annual, epiphytic, temperate, and subtropical genera, and comparatively weak (though still significantly positive) for tropical and woody (tree, other woody perennial) genera (Fig 2b).

Across families included in this analysis (100 families; 91.8% of species), the genus size -range size relationship was mixed (Fig 2c); 54 families have significant (p<0.05), positive slope coefficients (i.e. species in larger genera are more likely to be restricted to a single botanical country), five families have significant negative coefficients and 41 families had slope coefficients not significantly different from zero.

### Genus size and extinction risk

Species in big genera are less likely to have a global Red List assessment (Fig 3); for a monotypic genus, the probability of being assessed is 15.3% (95% CI: 14.4-16.2%), compared to 11.5% (CI: 11.2-11.7%) for a species in a 500-species genus (estimated marginal means).

**Figure 3.**
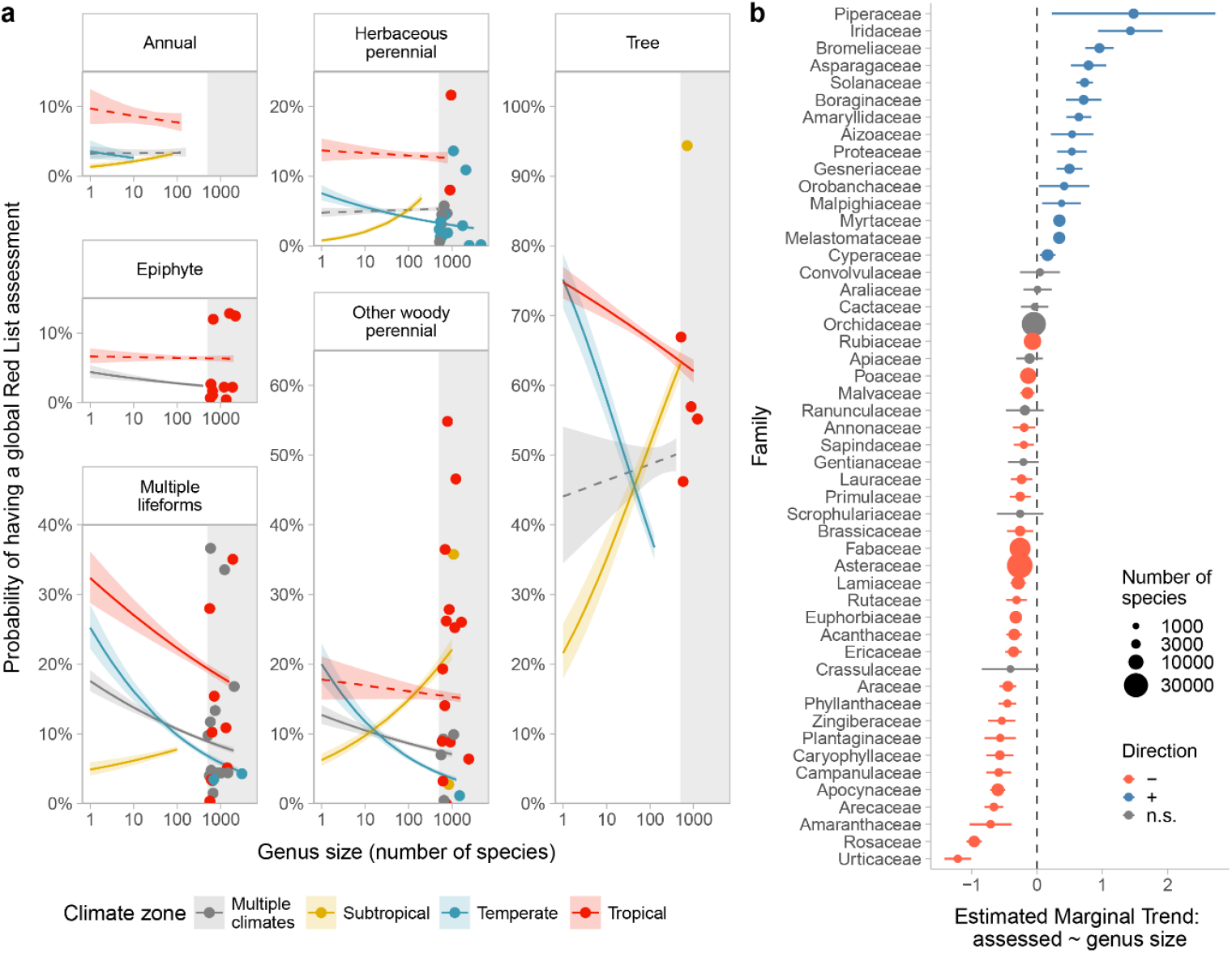
Genus size-assessment probability relationship. Panel a) shows estimated marginal means of assessment probability across genus size, climate and lifeform (note that climate and lifeform were aggregated at genus level). Upper limit of genus size values restricted to range of data for each covariate combination. Grey rectangle highlights the ‘zone of big genera’; points show the observed proportions of assessed species in big genera. Dashed lines show n.s. trends; solid lines show trends statistically different from zero. Panel b) shows the estimated marginal slope coefficients for family (the single-country endemic threshold; only largest 50 families shown); negative coefficients indicate that species in larger genera are less likely to have a Red List assessment. Units of coefficients on the logit scale. Error bars show 95% CIs of coefficient estimates; colour of points denotes significance and direction; size of points denotes number of species per family. Marginal means and trends estimated across cell weights of the non-focal terms in all panels; see Fig S1 for a graphical representation of how these are estimated from the model. For tabulated results with 95% CI values for marginal climate, lifeform, and family trends, see Table S4; for all trends (not averaged), see Table S5.

Marginal trends across lifeform and climate are generally consistent with the overall effect, except for subtropical genera – species in subtropical genera are more likely to be assessed if they are in a larger genus (Fig 3a). This result is likely driven by the near-comprehensive (94%) assessment of *Eucalyptus* (Myrtaceae), the only large subtropical tree genus, comprising 27.0% of species (across all lifeforms) in big subtropical genera. The impact of the Global Tree Assessment is clearly visible in our data, with trees having the highest assessment coverage of any covariate (Fig 3a).

Marginal trends across families are less consistent: 49 families have significant negative trends as expected, 19 have positive trends and 32 are not significantly different from zero (50 largest families shown in Fig 3b). Regardless of lifeform, species in tropical genera are always more likely to have been assessed than those from other climates (Fig. 3a).

Using Red List data, we found that when assessed, species in big genera are slightly more likely to be assessed as threatened (either Vulnerable, Endangered or Critically Endangered; Fig 4a). On average, an assessed species in a monotypic genus has a 32.6% (CI: 31.0-34.1%) chance of being assessed as threatened, compared to 37.5% (CI: 36.8-38.2%) if it is in a genus of 500 species (estimated marginal means). The effect is generally reversed in epiphytic and subtropical genera (Fig 4b), though this is likely driven by a small number of assessments, as species from epiphytic or subtropical genera tend to be less likely to have assessments (Fig 3). Across the 51 families with sufficient assessment coverage for analysis (Fig 4c), 13 have a significant positive relationship as expected, 9 have a significant negative relationship and 29 have relationships between genus size and probability of being assessed as threatened that are not significantly different from zero.

**Figure 4.**
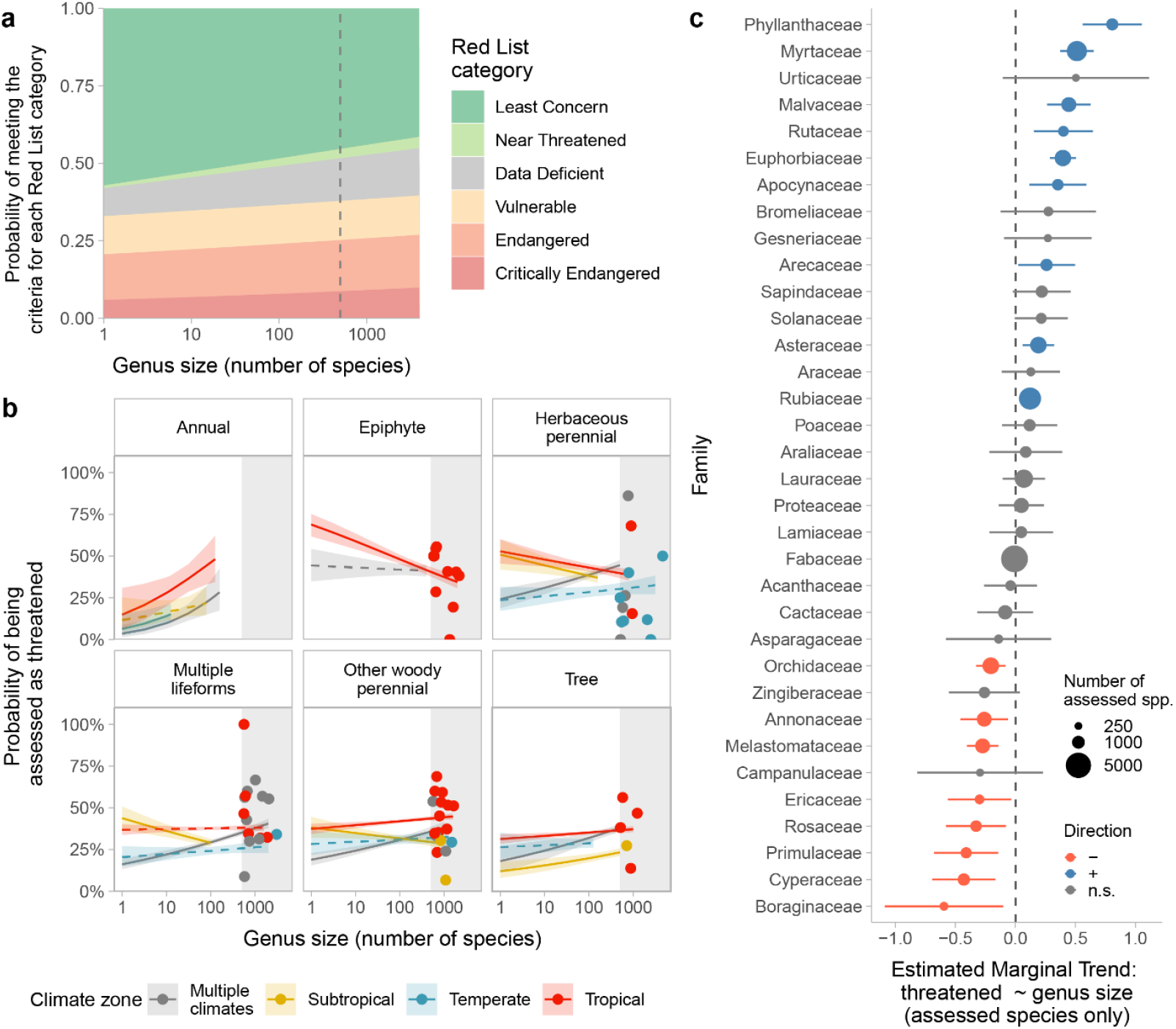
Genus size - documented extinction risk relationship (Red List assessments only). Panel a) shows estimated marginal means for the probability of meeting the criteria for each Red List category (note that categories are cumulative, e.g. all species assessed as Critically Endangered also meet the criteria for Endangered and Vulnerable. Panel b) shows estimated marginal means of being threatened (Vulnerable, Endangered or Critically Endangered) across genus size, climate and lifeform (note that climate and lifeform were aggregated at genus level). Upper limit of genus size values restricted to range of data for each covariate combination. Grey rectangle highlights the ‘zone of big genera’; points show the observed proportions of species assessed as threatened in big genera. Dashed lines show n.s. trends; solid lines show trends statistically different from zero. Panel c) shows the estimated marginal slope coefficients for family (only 34 of the largest 50 families shown; 16 of those families have <250 assessed species so were excluded from this analysis); positive coefficients indicate that species in larger genera are more likely to be assessed as threatened. Units of coefficients on the logit scale. Error bars show 95% CIs of coefficient estimates; colour of points denotes significance and direction; size of points denotes number of species per family. Marginal means and trends estimated across cell weights of the non-focal terms in all panels; see Fig S1 for a graphical representation of how these are estimated from the model. For tabulated results with 95% CI values for marginal climate, lifeform, and family trends, see Table S4; for all trends (not averaged), see Table S5.

Species in larger genera are also more likely to be categorised as Data Deficient (Fig. 5); the probability that a species in a monotypic genus is categorised as Data Deficient is 4.5% (95% CI:4.1-4.9%), compared to 9.1% (CI: 8.7-9.4%) in a genus with 500 species. The effect on Data Deficiency of being in a large genus is strongest for tropical and epiphytic species, but reversed for annual and subtropical species (i.e. species in larger genera are less likely to be assessed as Data Deficient), though these latter groups have, respectively, only 18 and 94 species (zero and seven in big genera) that have been assessed as Data Deficient. Data were insufficient to fit the full model including family; full model for Data Deficiency fitted with climate and lifeform covariates only.

**Figure 5.**
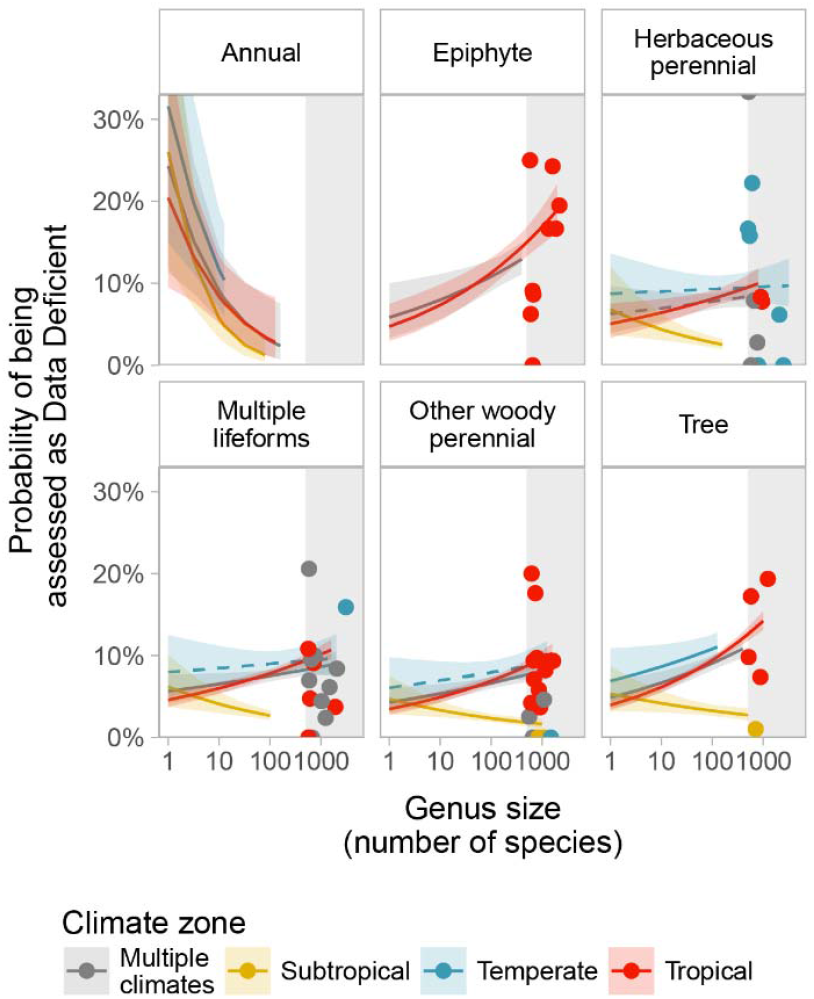
Genus size – Data Deficiency relationship (Red List assessments only). Estimated marginal means of being assessed as Data Deficient across genus size, climate and lifeform (climate and lifeform were aggregated at genus level). Upper limit of genus size values restricted to range of data for each covariate combination. Shaded ribbons show 95% CIs. Dashed lines show n.s. trends; solid lines show trends that are statistically different from zero. For tabulated results with 95% CI values for climate and lifeform trends, see Table S5.

When we extended our analysis of extinction risk to include all angiosperms, by including extinction risk predictions (from Bachman *et al*., 2024) for species lacking a Red List assessment, we found a stronger overall relationship between genus size and extinction risk (Fig 6): a species in a monotypic genus has a 30.4% (CI: 29.8-31.1%) chance of being threatened (assessed or predicted with high confidence), compared to 45.6% (CI: 45.3-45.9%) for a genus of 500 species (estimated marginal means).

**Figure 6.**
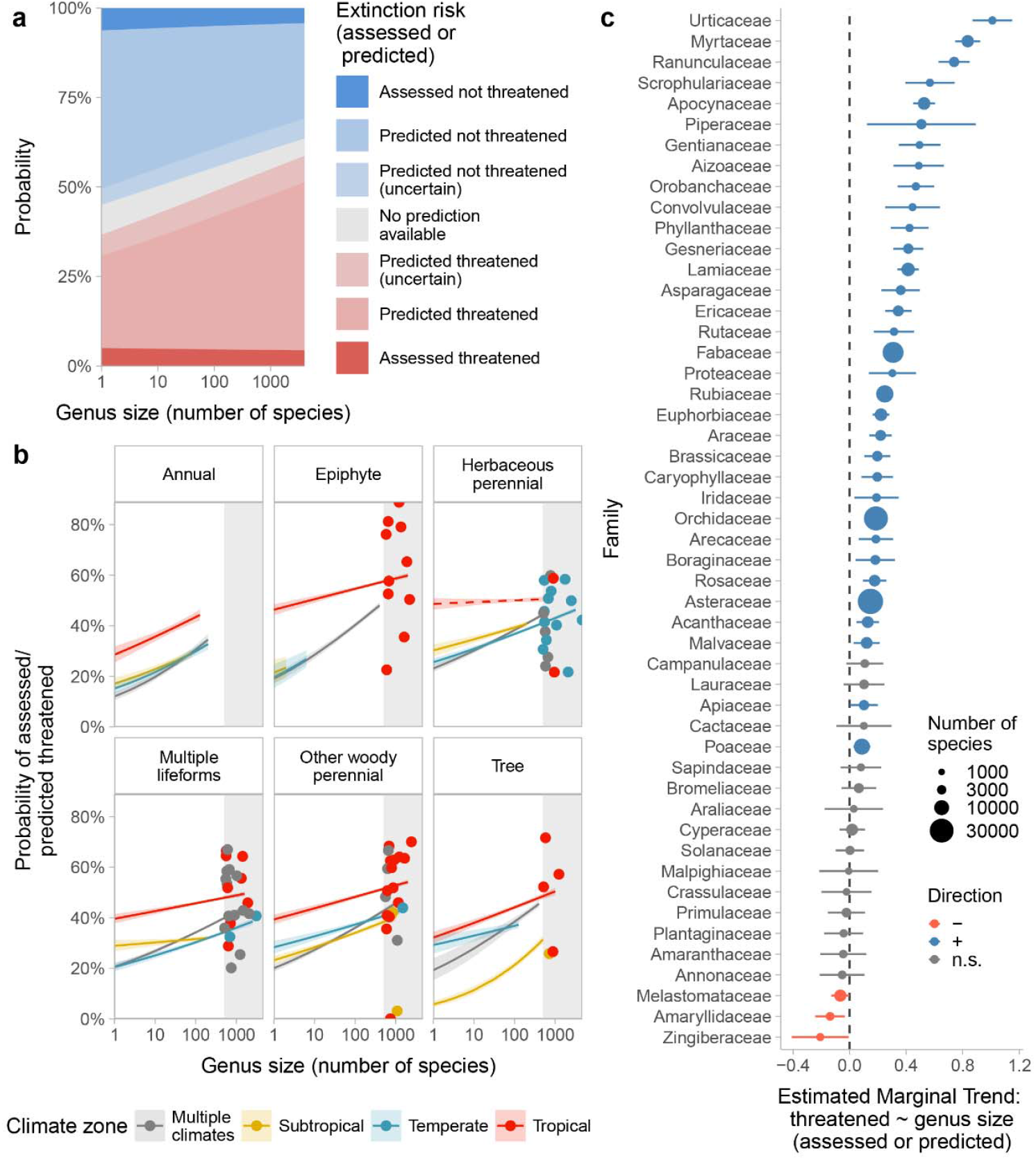
Genus size - extinction risk relationship (Red List assessments and extinction risk predictions). Panel a) shows estimated marginal means across all thresholds of estimated extinction risk. Panel b) shows estimated marginal means of being threatened (assessed or predicted with high confidence) across genus size, climate and lifeform (climate and lifeform were aggregated at genus level). Upper limit of genus size values restricted to range of data for each covariate combination. Grey rectangle highlights the ‘zone of big genera’; points show the proportions of species in big genera that are assessed or predicted to be threatened. Dashed lines show n.s. trends; solid lines show trends that are statistically different from zero. Panel c) shows the estimated marginal slope coefficients for family (only largest 50 families shown); positive coefficients indicate that species in larger genera are more likely threatened (either assessed or predicted). Units of coefficients on the logit scale. Error bars show 95% CIs of coefficient estimates; colour of points denotes significance and direction; size of points denotes number of species per family. Marginal means and trends estimated across cell weights of the non-focal terms in all panels; see Fig S1 for a graphical representation of how these are estimated from the model. For tabulated results with 95% CI values for marginal climate, lifeform, and family trends, see Table S4; for all trends (not averaged), see Table S5.

The relationship between genus size and predicted probability of being threatened is consistent across climatic zones and lifeforms (Fig 6b). It is strongest for annual, epiphytic and genera found in multiple climates, and comparatively weak (though still significantly positive) for tropical, subtropical and woody genera (Fig 6b). Across families (Fig 6c), the genus size – extinction risk relationship is broadly consistent, though statistical power is limited; of the 100 families included in this analysis, 56 have significant (p<0.05), positive slope coefficients (i.e. species in larger genera are more likely to be either assessed as or predicted to be threatened), five have significant negative coefficients and 39 have slope coefficients that are not significantly different from zero. When we compared estimates of the predicted proportions of threatened species based on assessments versus assessments plus predictions (Fig 7), we found evidence to suggest that current Red List data underestimate the proportion of species that are threatened with extinction in genera of eleven or more species. This underestimation is most severe for big genera.

**Figure 7.**
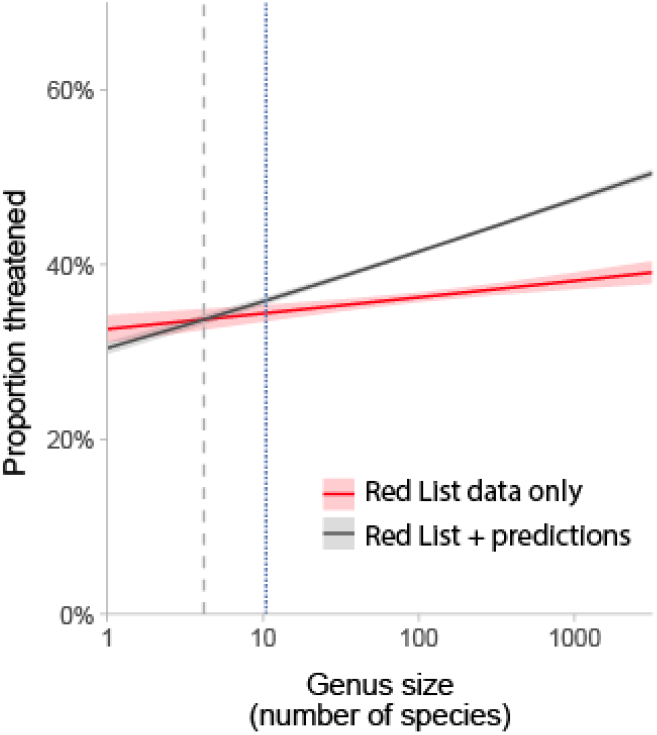
Comparison of the estimated marginal means for Red List data (red) and combined (Red List + predictions) data (grey) for the relationship between genus size and probability of extinction risk. Dotted lines show where the lines intercept (dashed grey; left) and where the 95% CIs diverge (dotted blue; right), indicating the change from over-to under-estimation of threat.

From the extinction risk predictions (Bachman *et al*., 2024), 37,072 species in big genera are estimated (with high confidence) to be threatened with extinction, but either lack a global Red List assessment (36,411) or have been assessed as Data Deficient (661) – together representing 10.9% of all angiosperm species. This number is conservative, as >13,000 species have been described or otherwise added to WCVP since the release used as the backbone for predictions. Of the species in big genera predicted to be threatened, 19,787 are in tropical genera (spanning 40 genera). We quantified the degree to which extinction risk is likely to be underestimated in each large genus by subtracting the Red-List-derived proportion of threatened species from that calculated using predictions (Table 1). Of the twenty big genera in which extinction risk is likely to be most underestimated on the Red List, six are tropical genera in the family Orchidaceae. The most extreme example is *Stelis* with <1% of species having a Red List assessment and none threatened, while 79% of species are predicted to be threatened.

**Table 1.**
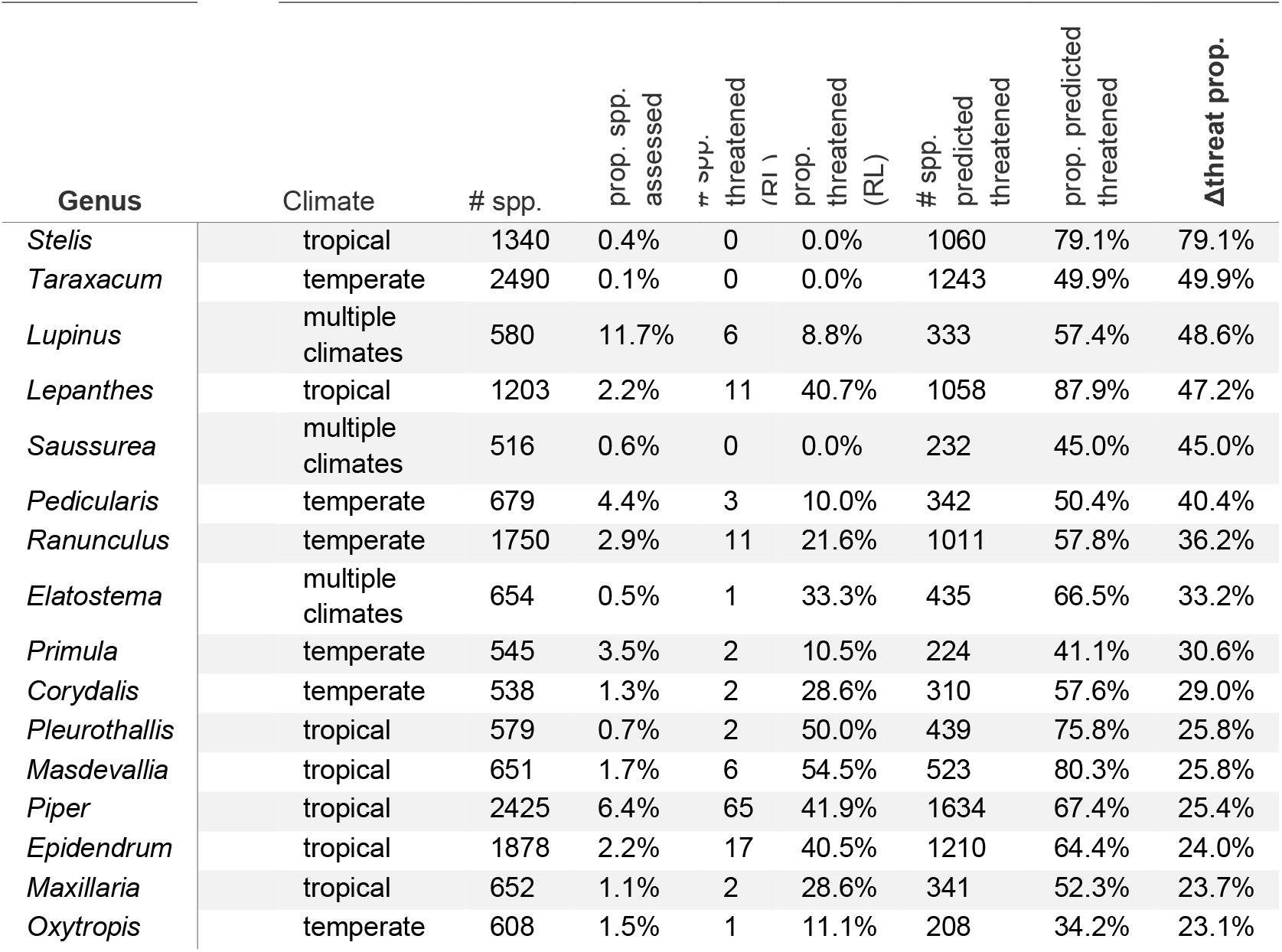
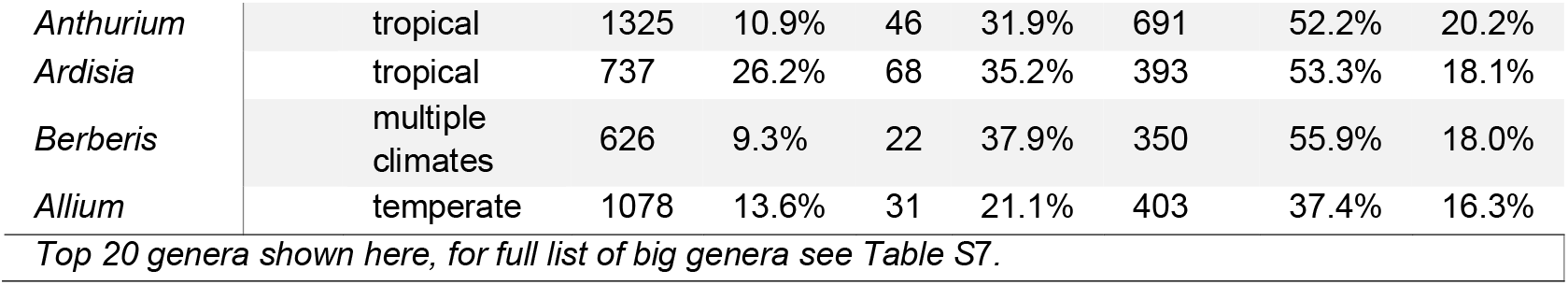
Numbers of species assessed as versus predicted to be threatened in big genera, arranged in descending order of underprediction (Δthreat prop.).

## Discussion

Consistent with our expectations, being in a larger genus is associated with having a smaller geographic range, being less likely to have a Red List assessment, and if assessed, being more likely to be either Data Deficient, or assigned to one of the threatened categories. The relationships between genus size, range size and extinction risk are broadly consistent across climatic zones, lifeforms, and families; as over 25% of plant species are in ‘big’ genera (≥500 species) and a further 35% are in moderately large genera (≥100 species), our findings are relevant to the majority of known plant species and have significant implications for conservation science, action and policy.

Our results support previous studies on the relationship between taxon size and range size in plants (in genera; Schwartz & Simberloff, 2001; Lozano & Schwartz, 2005; Leão *et al*., 2020; and families; Fu *et al*., 2022). However, the relationship between genus and range size is not universal – in five families, species in larger genera are less likely to be restricted to a single botanical country, contrary to expectations (Table S4). These families span seven orders (both eudicots and monocots), vary in climate, lifeform and range sizes, and three contain at least one big genus, so we found no clear pattern to these outliers. Similarly, we found groups (discussed below) for which the general pattern of higher extinction risk in larger genera is reversed, where species in larger genera are *less* likely to be either assessed as or predicted to be threatened (Figs 3-4; Tables S4-S6).

Results based on published assessments were less consistent than those based on predictions, likely due to the very low assessment coverage of many big genera. For example, in tropical epiphytes, where seven of the ten big genera have fewer than 50 published assessments (representing <3% of species in these seven genera), assessments show an apparent negative relationship between genus size and extinction risk (Fig 4), but the model including predictions showed the expected positive trend (Fig 5). There are five families with a negative relationship between genus size and extinction risk in prediction-based models. For one of these, Melastomataceae, this may be linked to range, as it was one of the five families where species in larger genera tended to have larger ranges (Table S4), but for the other four the reasons for this result are unclear; differences in both range size and extinction risk may be linked to the ‘different ways to be a big genus’ (an outstanding knowledge gap noted by Moonlight *et al*., 2024). It is also possible that there are biases in the predictions that affect our results – for example, the phylogenetic eigenvectors used as predictors by Bachman *et al*. (2024) make it more likely that predictions match the Red List-derived estimates of extinction risk within a genus (though this effect seems to be weak; Table 1). These exceptions do not in any way diminish the need for species in big genera to be prioritised for conservation assessment, whether to formally document their predicted extinction risk, address the gaps in both knowledge and data for big genera or to better understand the drivers of extinction risk across taxon size.

The mechanisms underpinning the link between genus and range sizes may be related to evolutionary diversification patterns. More rapidly diversifying plant genera tend to have smaller ranges and higher extinction risk (Davies *et al*., 2011; Leão *et al*., 2020; Tanentzap *et al*., 2020) though conifers showed the opposite trend (Tanentzap *et al*., 2020). In animals, genus-level diversification rate appears to be a strong correlate of both range size and extinction risk (though there is significant variation across clades; Greenberg *et al*., 2021), further supporting the hypothesis that rapid speciation tends to produce range-restricted species which are inherently more vulnerable to extinction (Davies *et al*., 2011; Leão *et al*., 2020). Our findings regarding higher extinction risk of species in larger genera are in contrast to earlier reports for both plants (Vamosi & Wilson, 2008) and animals (Purvis *et al*., 2000) that smaller – especially monotypic – taxa tend to have higher levels of extinction risk.

Our results complement recent studies characterising biases in the representation of plants on the Red List and addressing how these can be quantified and corrected (Nic Lughadha *et al*., 2020) or circumvented (Bachman *et al*., 2024) in future analyses to refine understanding of global and regional patterns of plant extinction risk. For example, eight of the 20 most under-estimated genera (when overall extinction risk is estimated using the Red List; Table 1) are Orchidaceae (6 genera) or Asteraceae (2 genera). These two largest plant families encompass c. 65,000 species and are recognised as being among the most under-represented on the Red List (Nic Lughadha *et al*., 2020).

Strikingly, the six orchid genera highlighted here for their underestimated threat levels are all tropical, consistent with our original expectation that under-representation of big genera in assessment processes might be greatest in the tropics. However, contrary to our expectations, we found that species in tropical genera, across all lifeforms and independent of genus size, were more likely to have been assessed than species of other climates (Fig 3a). This encouraging result suggests significant progress towards addressing the plant Red Listing deficit which was so evident across many areas of the tropics as recently as a decade ago (Bachman *et al*., 2019). Much of this progress is attributable to the Global Tree Assessment, and the high assessment coverage achieved for trees explains the relatively weak effect of genus size on assessment probability in tree-dominated genera. Similarly concerted and large-scale endeavours are urgently needed for other plants, and our findings suggest that an explicit focus on comprehensive assessments of selected big genera could be an important and worthwhile approach to start to redress the effects of neglect or avoidance of these groups over decades.

We are in no way suggesting that large genera be split, or that there should be an upper limit on the size of genera. The size of genera is related to both evolutionary and human processes (e.g. Bentham’s preference for fewer, larger genera; Humphreys & Linder, 2009), and the highly skewed pattern of genus size frequency, with many small and relatively few very large genera, is common across the tree of life (Strand & Panova, 2015; Sigwart *et al*., 2018). Although ultimately arbitrary, there is a broad consensus that ‘good’ genera (monophyletic, stable and diagnosable) are useful divisions (Humphreys & Linder, 2009; Vences *et al*., 2013; Strand & Panova, 2015; Muñoz-Rodríguez *et al*., 2023), so big plant genera (and their problems) are unlikely to be split out of existence (Moonlight *et al*., 2024). However, regardless of the basis of genus size, our results show that by treating genus size as an attribute of species, we can quantify the tangible conservation implications of being a member of a big genus.

We strongly encourage greater focus on and resourcing of the further study of species in big plant genera in taxonomic, ecological and conservation contexts. Although big genera continue to represent an immense amount of work to monograph *in toto*, molecular data can be used to divide large taxa into more manageable subclades; as in *Eugenia* or *Myrcia* (Mazine *et al*., 2014; Lucas *et al*., 2018). An alternative may be the ‘foundation monograph’ concept (Scotland & Wood, 2012) – a practical guide to this approach (based on the recent monograph of *Ipomoea*; Muñoz-Rodríguez *et al*., 2019) is detailed in Muñoz-Rodríguez *et al*. (2024). We echo earlier recommendations for increased collaboration across conservation and taxonomy (Nic Lughadha *et al*., 2019; Brown *et al*., 2023a). The success of any new initiative focused on assessing big genera will depend on a clear-eyed, evidence-based understanding of the challenges likely to be faced; among these is the higher proportion of Data Deficient species in genera which lack recent monographs (Grace *et al*., 2021) and which are likely to include unusually large proportions of species not yet described (Moonlight *et al*., 2024). We found that species from big genera were at least twice as likely to be categorised as Data Deficient as those in monotypic genera, a result consistent with our expectations. However, this proportion remained relatively low overall (<10%, Fig 5) and could be further minimised by effective collaboration between taxonomists, assessors and other local experts to assemble the most complete information possible for species’ assessments. Where assessors identify species likely to be Data Deficient, this can feed back into research prioritisation (e.g. targeted fieldwork or other data collection).

Similarly, taxonomists can provide input on the likelihood of species re-circumscriptions, facilitating effective prioritisation of assessments in big genera – thus, a shared pipeline of activity between taxonomists and assessors can allow extinction risk assessments of species in speciose genera to be undertaken before a genus is fully monographed (Nic Lughadha *et al*., 2019).

Species in big genera are undoubtedly conservation priorities; they are under-studied, under-assessed, and their extinction risk has been underestimated. Over a quarter of angiosperm species diversity is in big genera, but there is a systematic shortfall in extinction risk data for these species. Moreover, we found that this knowledge gap is likely to obscure, at least in part, the greater extinction risk faced by species in big genera than by other angiosperms. While working with speciose taxa presents ongoing practical challenges, an urgent increase in focus on studying, assessing and conserving species in big genera is vital to safeguard the future of angiosperm diversity.

